# Uniform pre-processing of bacterial single-cell RNA-seq

**DOI:** 10.64898/2025.12.04.692398

**Authors:** Conrad Oakes, Vera Beilinson, Margaret J. McFall-Ngai, Lior Pachter

**Affiliations:** Division of Biology and Biological Engineering, California Institute of Technology, Pasadena, CA, 91125, USA; Division of Biosphere Sciences and Engineering, Carnegie Institution for Science, Pasadena, CA, 91125, USA; Department of Computing and Mathematical Sciences, California Institute of Technology, Pasadena, CA, 91125, USA

**Author notes:** Correspondence: Lior Pachter.

## Abstract

Bacteria are highly heterogeneous, even under controlled conditions, making single-cell RNA sequencing (scRNA-seq) essential for studying microbial diversity and symbiosis. Since its first application in 2015, bacterial scRNA-seq has expanded, but different assays depend on distinct, custom, in-house preprocessing making it difficult to analyze data as part of a unified workflow. The kallisto-bustools suite of tools has enabled uniform pre-processing of eukaryotic scRNA-seq while also reducing time and resource demands for pre-processing, but is not optimized for bacterial scRNA-seq. We adapt kallisto-bustools to be suitable for reads generated from operons, as well as for a much shorter gene length distribution, and show that it can efficiently and accurately quantify bacterial scRNA-seq. Our work provides a scalable foundation for uniform pre-processing and analysis of microbial single-cell transcriptomics.

## Introduction

Single-cell RNA sequencing has transformed eukaryotic cell analysis by allowing an improved understanding of diversity at the individual-cell level. However, implementation of single-cell transcriptomics in bacteria has been limited due to a variety of biological and technical challenges, most significantly the lack of polyA tails. Recently, several methods have been developed that overcome these challenges, allowing untargeted sequencing of a bacterial RNA profile. With the growing recognition of the central role of host-bacteria interactions, bacterial single-cell transcriptomic datasets are poised to greatly increase in number (Figure 1).

**Fig. 1.**
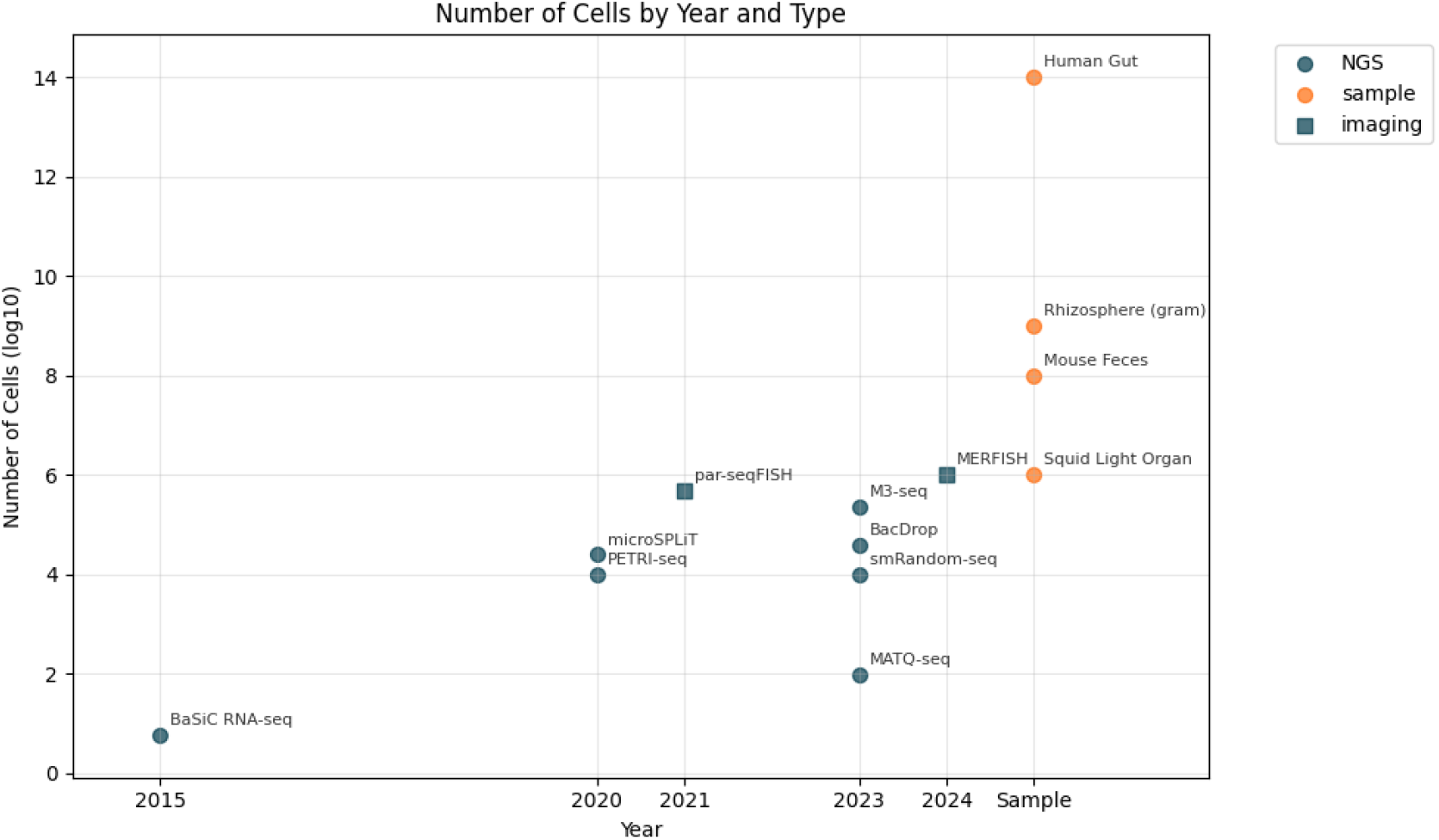
The number of cells analyzed was based on the values reported in each original method paper. For imaging-based approaches, cell numbers were determined by the practical limits of imaging throughput. Imaging-based methods are inherently targeted, requiring prior knowledge of the genes of interest to design appropriate probes. Biological samples (orange) do not have defined cell number; therefore, approximate values were used.

The analysis of bacterial single-cell RNA-seq data can be challenging, starting with the pre-processing of reads for gene quantification. Bacterial genomes are frequently poorly annotated (1), leading to quantification errors due to unknown operons, and existing eukaryotic-optimized tools may not perform as expected due to incompatible parameter choices. In eukaryotes, genes are typically separated from each other by non-coding regions and are transcribed individually. In that context, a sequencing read is usually assigned to a single gene. Reads that map equally well to multiple genes are flagged as “multimapped” or “ambiguous” to avoid over-counting and artificially inflating expression levels. However, bacteria do not follow this one-gene-per-read rule and can frequently produce polycistronic transcripts from operons, thus sequencing reads can span multiple adjacent genes. Thus, current scRNA-seq pre-processing tools not only miscount genes but can also miss entire gene expression profiles.

We show that the kallisto-bustools (kb-python) suite of tools (2–4), which provide accurate and efficient preprocessing solutions for eukaryotic bulk and single-cell RNA-seq datasets via pseudoalignment, can be optimized for bacterial pre-processing and can recapitulate the results of other tools in a small fraction of the time. Moreover, we show that with kb, quantification can be performed for data from multiple different bacterial scRNA-seq technologies (specifically PETRI-seq (5), MATQ-seq (6), and BacDrop (7)) and species (E coli, Salmonella Typhimurium, and K. pneumoniae), making possible uniform pre-processing of data generated by different laboratories for joint downstream analysis.

## Results

### PETRI-seq

We analyzed E. coli data from (5) and found that quantification with kb-python proceeded at a rate of 1.5 billion reads per hour, a 50-fold improvement over the 0.03 billion reads per hour of the default workflow from (5).

After pre-processing the data with kb-python, we found that 54.1 percent of reads pseudoaligned and yielded 16,384 total unique molecular identifiers (UMIs). We were also able to recapitulate the results of Figure 2f from (5) with the count matrices generated by kb-python (Figure 2).

**Fig. 2.**
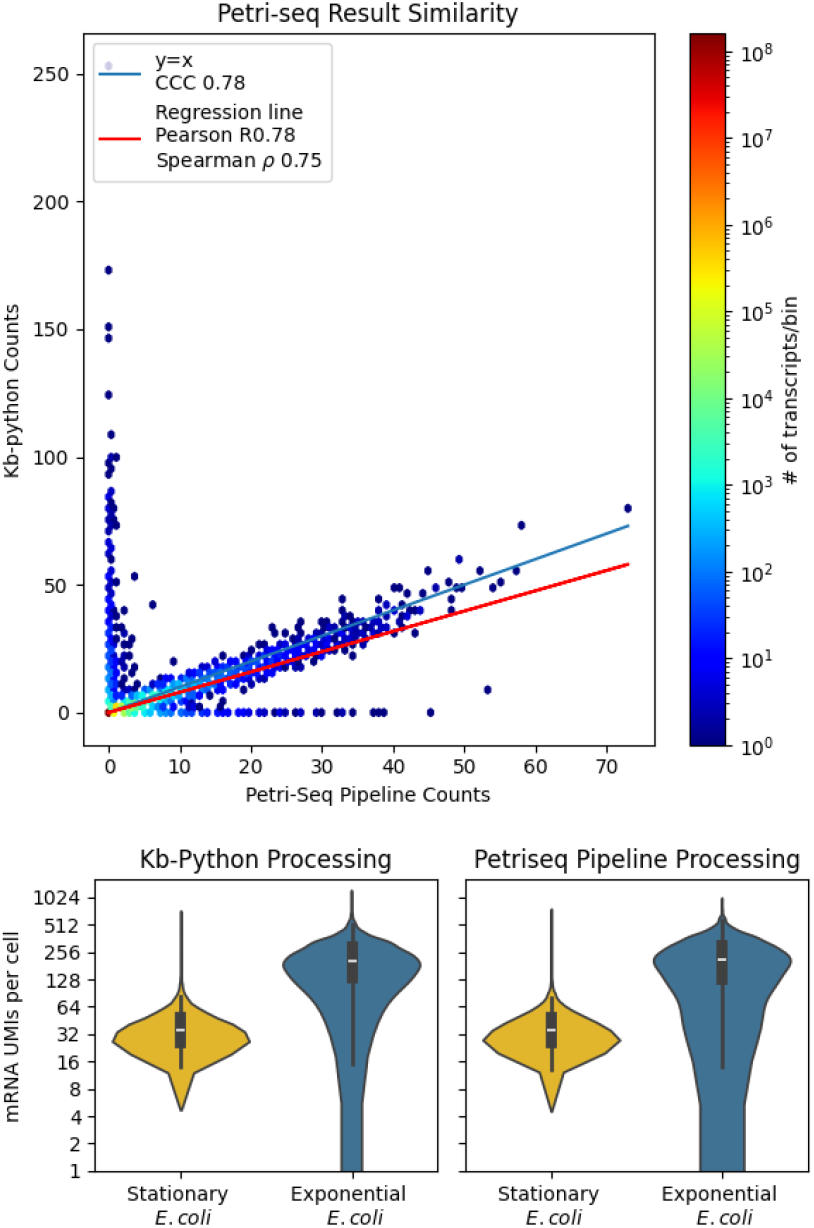
Quantification of the results of using kb-python on PETRI-seq compared to the standard workflow.

Applying a standard single cell workflow to the dataset, we used a filtering cutoff of 100 reads per cell based on the knee plot (Figure 6a). Exploring the mean-variance relationship, we found the data to be over-dispersed with a negative binomial distribution providing a reasnable fit (Figure 6b). Cells with more reads displayed more genes, but the library did not appear to have been saturated. Very few cells showed a high percentage of ribosomal genes expressed. Gene expression was then depth normalized and logged. The PCA representation of the data showed a clear separation between exponential and stationary phase cells (Figure 6e), and Leiden clustering showed that the stationary phase populations were much more heterogeneous than the exponential phase population. Upon inspection of the principal components, the main genes driving these differences appeared to be ribosomal. We further investigated this result by performing differential gene expression analysis using the edgePython program (8, 9), which showed the top differential genes to be related to metabolic processes (Figure 9).

We also explored the importance of operon identification in this dataset, which was made possible thanks to *E. coli* having a more well-defined genome than other bacteria species. We found increased correlation between genes within operons than those on different operons (Figure 10).

### MATQ-seq

We next analyzed Salmonella Typhimurium data from (6). Pre-processing with kb-python proceeded at a rate of 2.3 billion reads per hour, a more than 20-fold improvement over the 0.1 billion reads per hour of the default workflow of (6).

After pre-processing with kb-python, we found that 21 percent of reads pseudoaligned, yielding 321,945 “UMIs.” Our results showed high concordance with those reported using the workflow of (6) (Figure 3). Total reads were significantly higher than the other methods tested, presumably due to the lack of UMIs preventing collapsing duplicates. Exploring the mean-variance relationship, we found the data to be over-dispersed consistent with a negative binomial distribution (Figure 7b). There was no strong correlation between total number of reads and number of genes found (Figure 7c). Very few cells showed a high percentage of ribosomal genes expressed. After depth normalization and application of the logarithm transform, the 2-D PCA projection of the data showed some separation between different phases of cells in the expected order. Leiden derived clusters also aligned with the expected phases (Figure 6f).

**Fig. 3.**
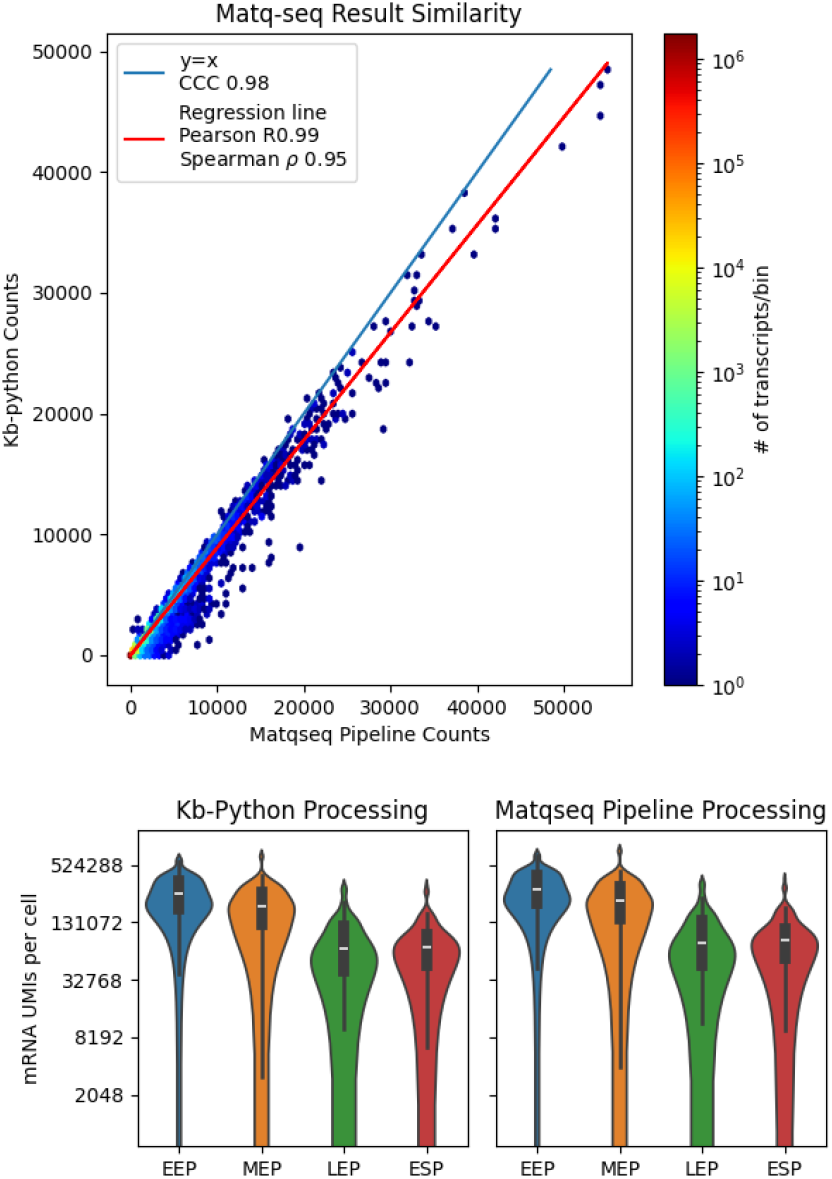
Quantification of the results of using kb-python on MATQ-seq compared to the standard workflow.

### BacDrop

Finally, we analyzed K. pneumoniae data from (7). Pre-processing with kb-python proceeded at a rate of 142 million reads per hour, a six-fold improvement over the 22 million reads per hour of the default workflow of (7).

After pre-processing with kb-python, we found that 50.1 percent of reads pseudoaligned, yielding 65,536 UMIs. Our results showed high concordance with those reported using the workflow of (7) (Figure 4). An obvious knee did not appear in the standard kneeplot (Figure 8a). Notably, the dataset had many more empty or near-empty cells compared to the other technologies. Exploring the mean-variance relationship, we found the data to be consistent with a negative binomial distribution (Figure 8b). Cells with more reads displayed more genes, but the library did not appear to have saturated. Several cells showed a high percentage of ribosomal genes expressed (Figure 8d). After depth normalization and application of the logarithm transform, the PCA representation and Leiden clustering show significant amoutns of heterogeneity, but there was no pre-defined metadata to explore the dataset, as was possible with the other datasets analyzed.

**Fig. 4.**
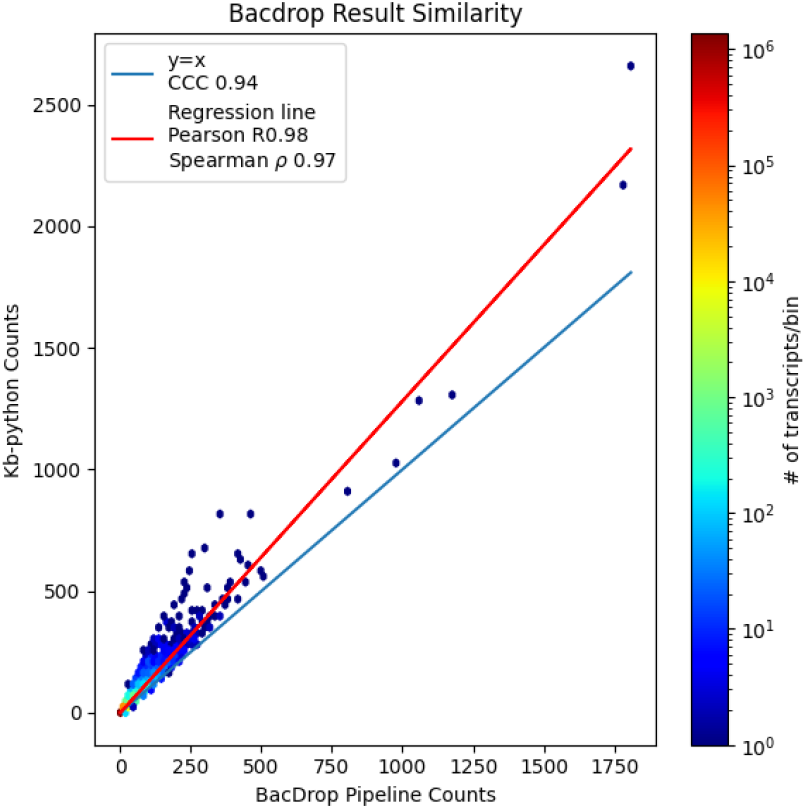
Quantification of the results of using kb-python on BacDrop compared to the standard workflow.

**Fig. 5.**
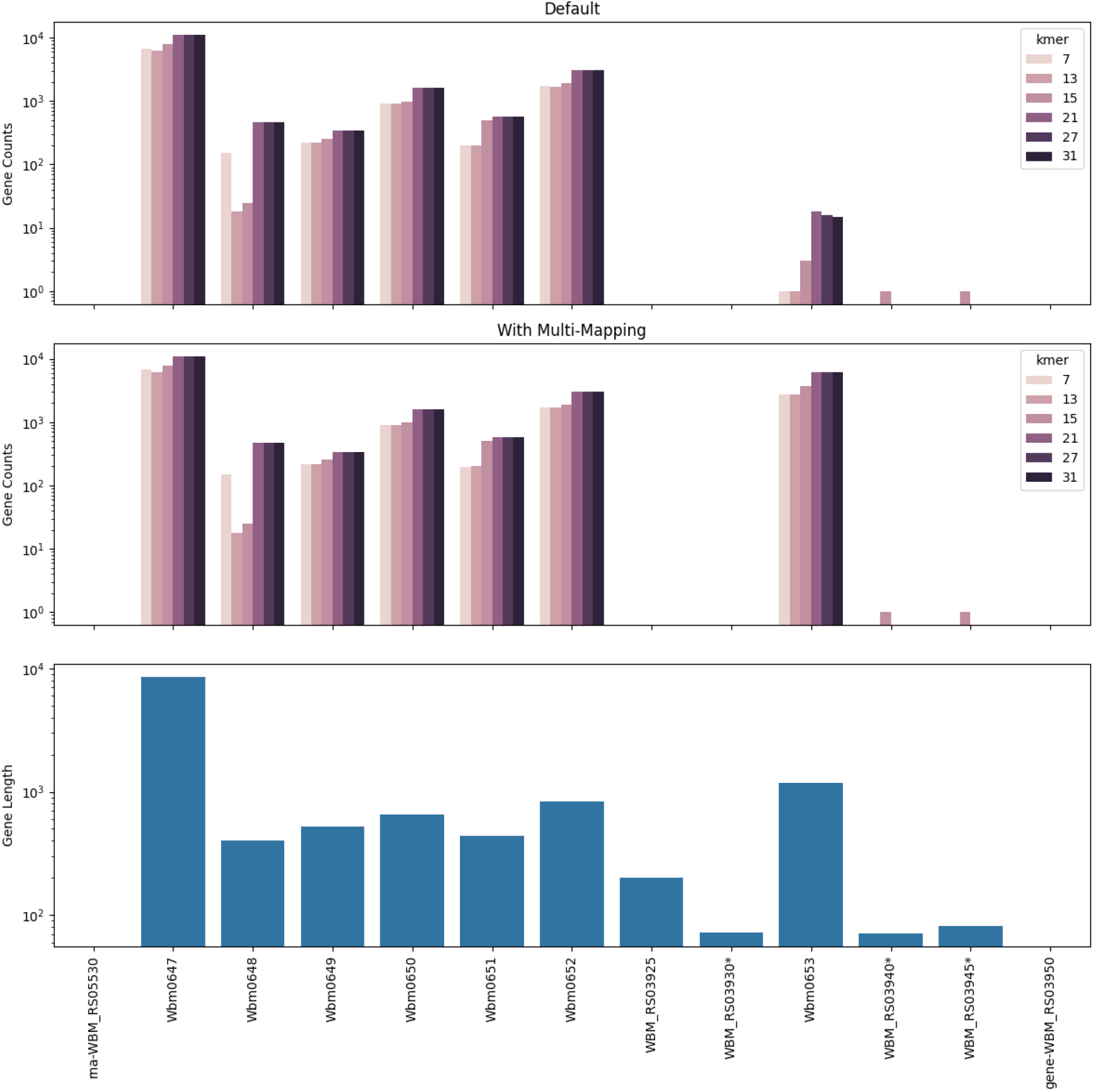
Quantification of the results of using kb-python on bulk RNAseq.

**Fig. 6.**
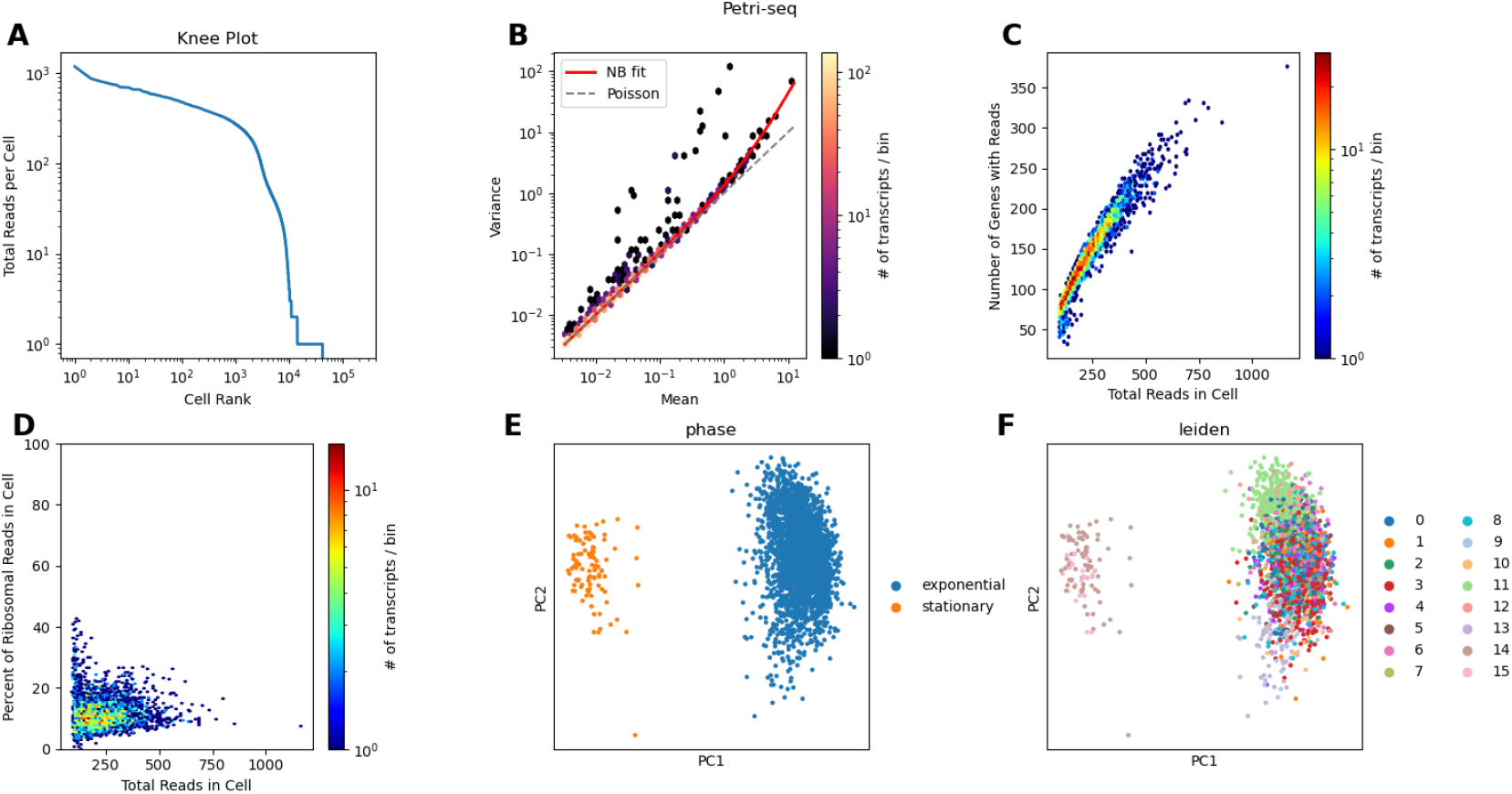
QC results on Petri-seq data.

**Fig. 7.**
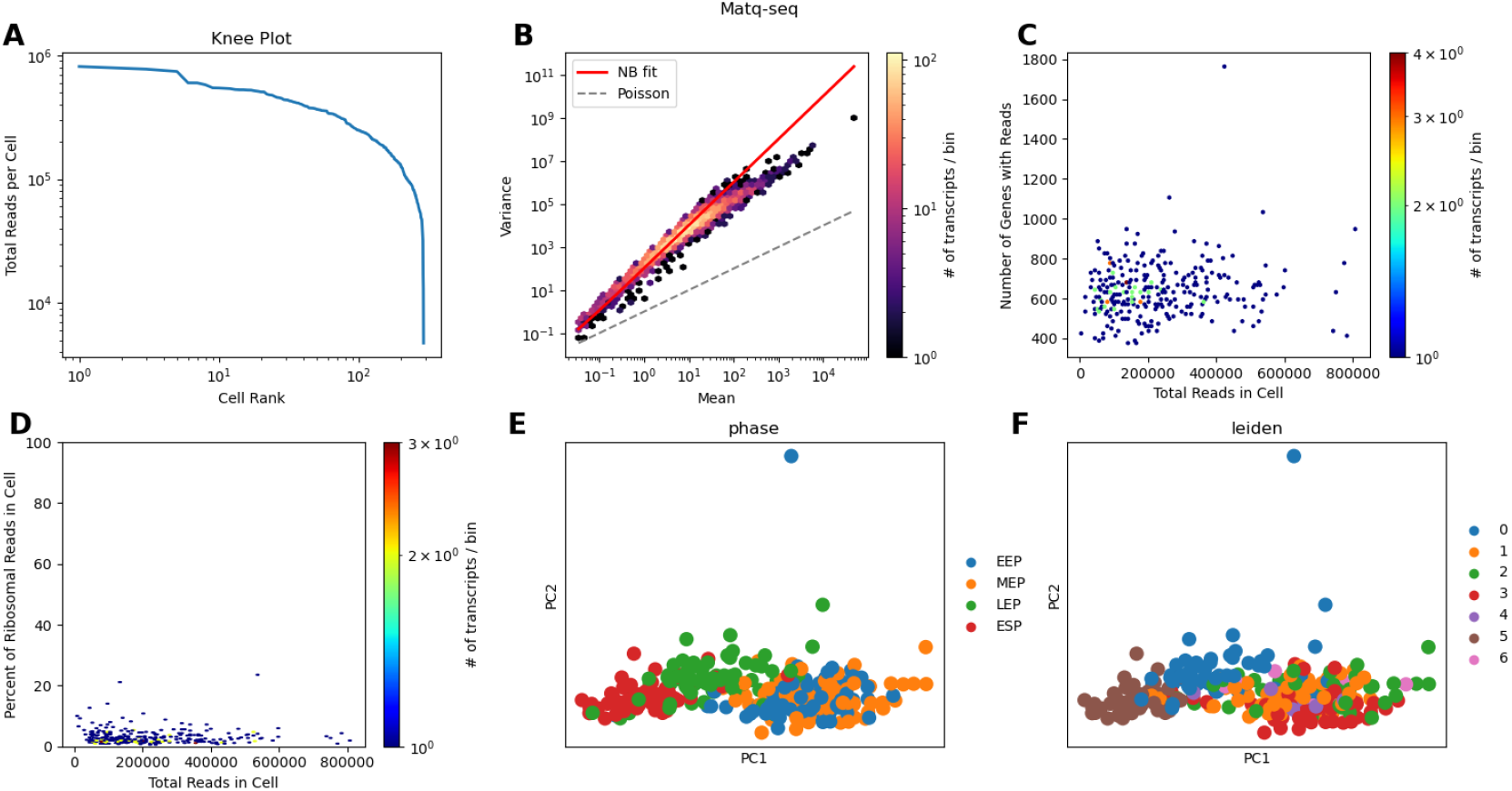
QC results on Matq-seq data.

**Fig. 8.**
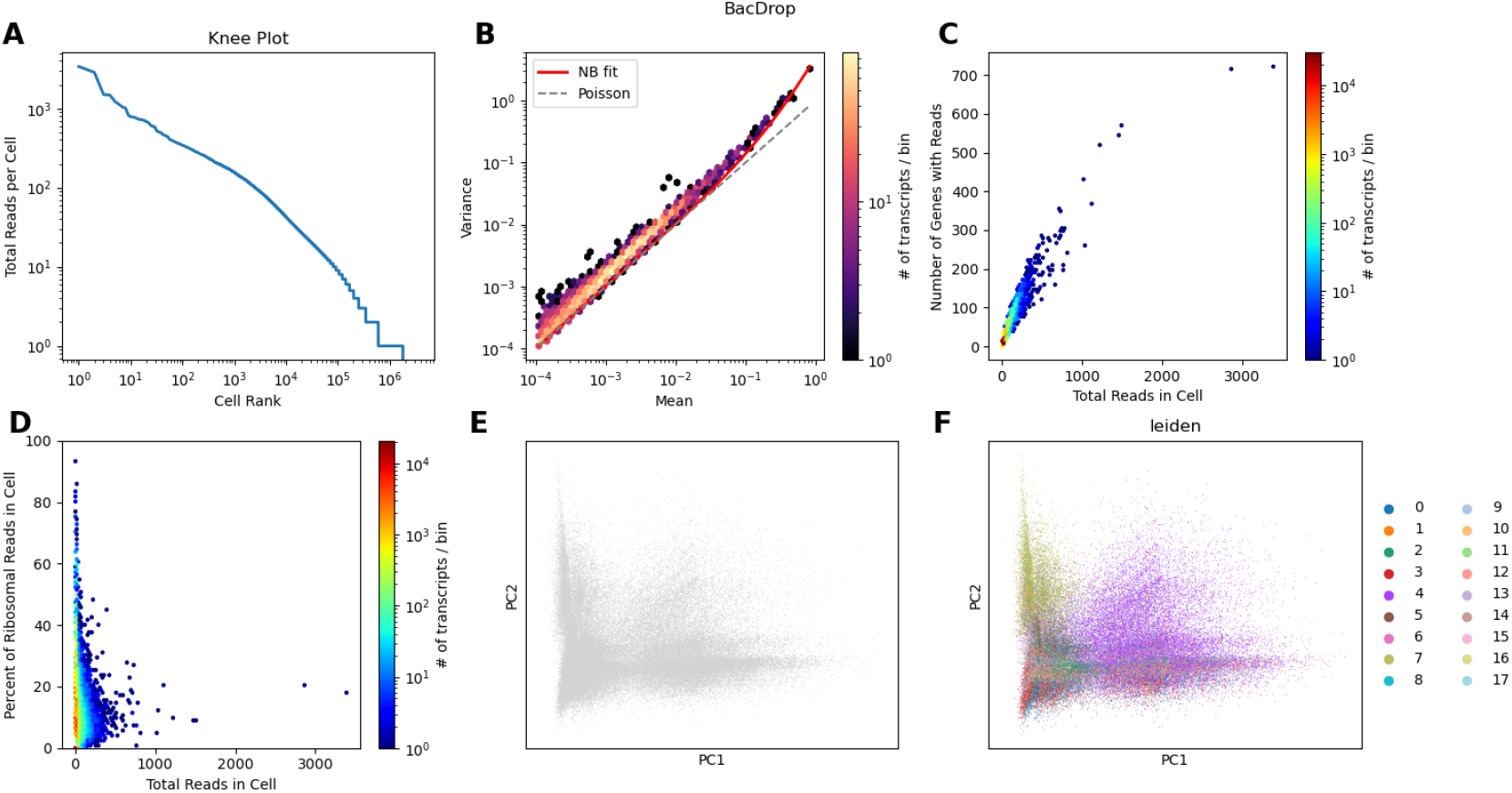
QC results on Bacdrop data.

**Fig. 9.**
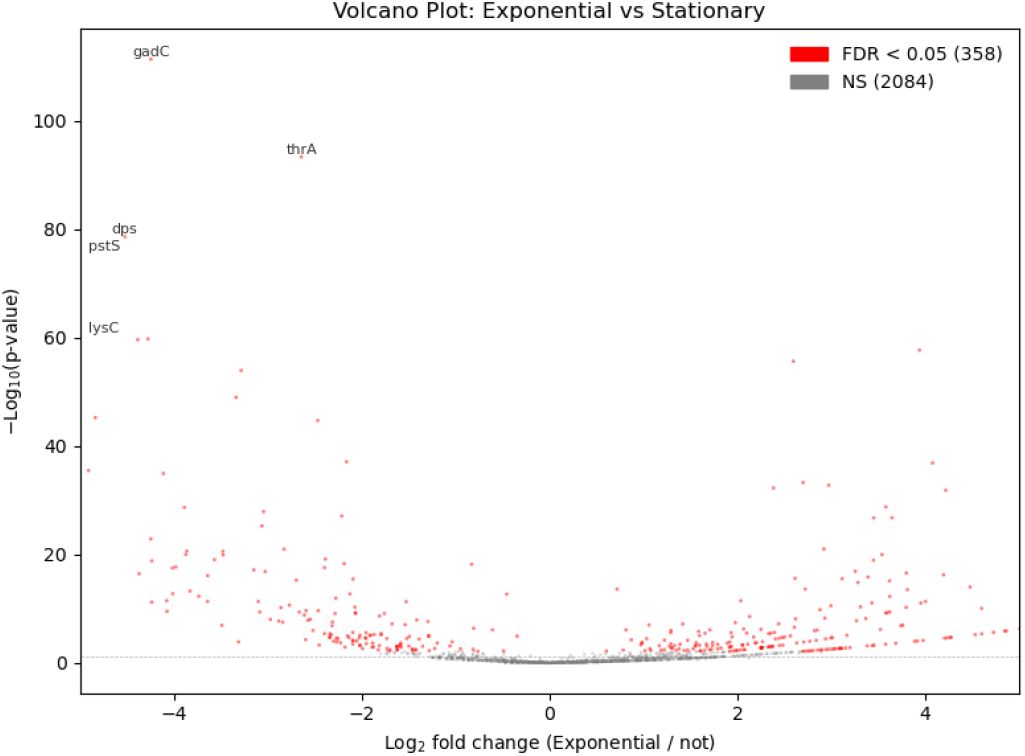
Volcano plot of DGE analysis of the Petriseq dataset.

**Fig. 10.**
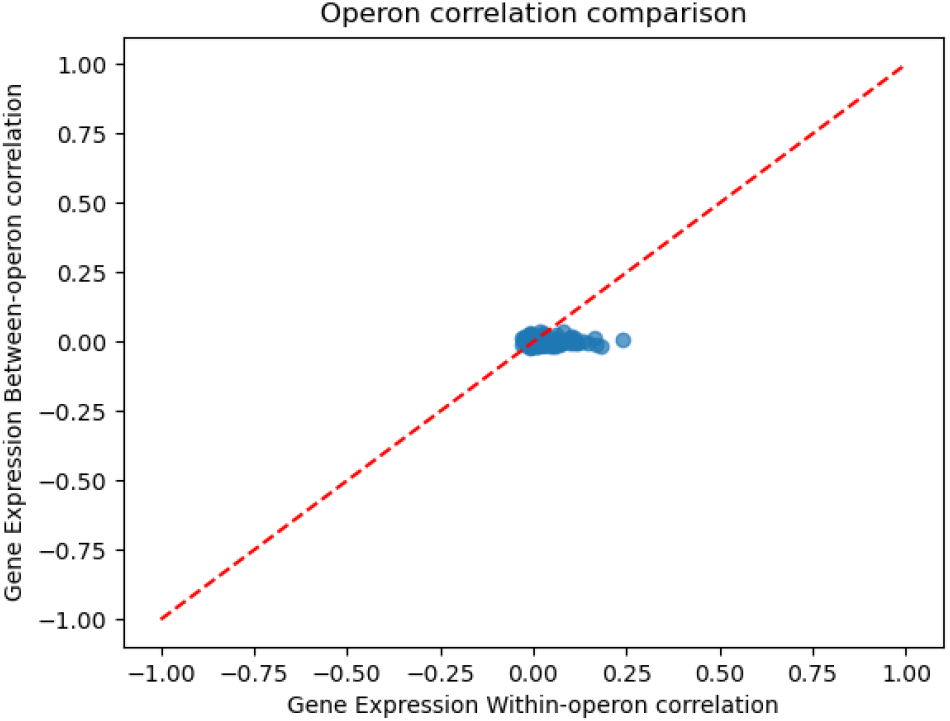
Correlation between gene expression within an operon and between other operons in the Petriseq dataset.

### k-mer selection and multimapping

Pseudoalignment has previously been reported to perform poorly on bacterial RNA-seq due to the removal of multi-mapped reads and missing small genes (10). To ascertain whether that is the case with kb, we assessed performance on a bulk RNAseq dataset from (10). We found that the kallisto built-in multimapping can recover expression that was previously reported as missing (Figure 5). Specifically, we examined the wBm stranded RNA-seq data set and found detection of more reasonable numbers for the gene Wbm0653. However, how best to make use of this feature will be specific to the exact organism/experiment in question, and thus is more a part of the downstream analysis of a potential user.

## Discussion

Our optimized pre-processing workflow demonstrates that kb-python can be effectively adapted for bacterial RNA-seq, making pre-processing highly efficient and accurate. Our kallisto-bustools workflow achieved 100-200X faster preprocessing times than possible with the tools accompanying the single-cell assays, which use traditional alignment-based approaches. Applying kb-python to two different bacterial scRNA-seq datasets, we were able to recapitulate the results from the original methods. Since viral transcripts have similar structure to bacteria, this optimized pre-processing workflow can be used for viral RNA-seq datasets as well.

The advantage of greatly reduced processing time with kallisto-bustools as compared with other tools is important, as future biologically relevant datasets are expected to be several orders of magnitude larger (Figure 1). Thus, our tool may help establish workflows that can be robust to the expansion of incoming data in the future (Table 1). Moreover, our approach should be amenable to other single-cell bacterial assays (7)

**Table 1.**
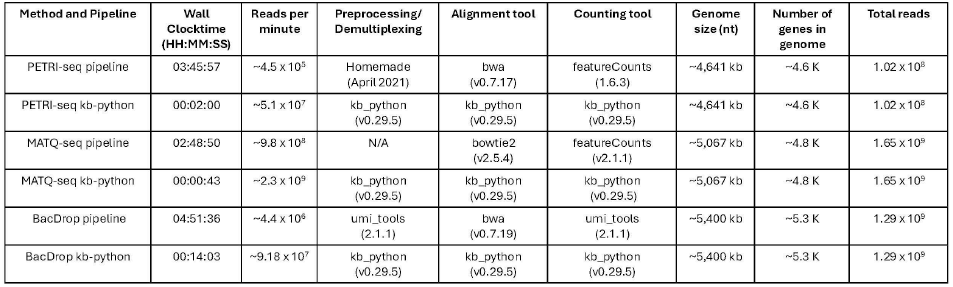
Benchmarking of the efficiency of kb-python compared to the workflow provided by the creators of each method (including tool version). Benchmarking was performed on an exclusive HPC compute node using scratch space to minimize interference from concert jobs. All workflows were run using 10 CPU cores and 50 GB of memory; scripts provided by method creators where modified for this by setting thread and memory threads on genome aligners and counting tools. Time will vary depending on machine, cores, and memory used.

One limitation of our method that must be considered is its robustness in the presence of contaminants. Single-cell bacterial RNA-seq can include unwanted bacterial organisms in the samples, and while kb-python can make use of indices that include other organisms, performance may be degraded. Unfortunately, the high similarity often seen between bacterial genomes can affect negatively pseudoalignment. This problem will become even more pronounced if the source of the contaminant is unknown.

One advantage of building a bacterial single-cell RNA-seq pre-processing workflow using kb-python is that it facilitates the use of other tools comprising the kallisto-bustools suite. For example, kallisto can be used to identify RdRP encoding viruses (11), making it possible to perform viral analyses alongside study of bacterial transcriptomics. Future expansion of the viral database could make it more well-suited to investigation of bacteriophages.

## Data and code availability

All the code to download data and generate the results of the paper is available at https://github.com/pachterlab/OBP_2025. The repository includes notebooks that can be directly run in Google Colaboratory. The kallisto program is available at https://github.com/pachterlab/kallisto. Documentation is at https://kallisto.readthedocs.io/.

## Acknowledgments

This project was funded by the Gordon and Betty Moore Foundation via grant GBMF12342 to M McFall-Ngai and by the NIH via grant NIH AI050661 to M McFall-Ngai and EG Ruby. We thank members of the McFall-Ngai, Ruby, and Pachter labs for helpful suggestions and feedback throughout the project.

## Methods

### Acquisition and preprocessing of PETRI-seq data

Raw RNA-seq reads from (5) were obtained from GSE141018; specifically GSM4489500/SRR11584024 for PETRI-seq Experiment 2.01. Reference genome *Escherichia coli* MB165 was used.

Files were passed into the kallisto/bustools workflow. For details and scripts on the workflow, see https://github.com/pachterlab/OBP_2025. The resulting count matrix was then loaded into scanpy for basic filtering.

Reads were also passed into the PETRI-seq workflow following code from https://tavazoielab.c2b2.columbia.edu/PETRI-seq/ using 40000 barcodes. `featureCounts_directional_5.py` was modified to count on the gene level instead of operon level (feature-Counts -t ‘gene’).

When timing was calculated for the PETRI-seq workflow, genome indexing, quality control with FastQC, and other pre-processing steps, were not included. For kb-python reference index generation was not included.

### Acquisition and preprocessing of MATQ-seq data

Raw RNA-seq reads from (6) were obtained from https://www.ncbi.nlm.nih.gov/geo/query/acc.cgi?acc=GSE218632. Reference genome https://www.ncbi.nlm.nih.gov/datasets/genome/GCF_000210855.2/ *Salmonella typhimurium* SL13444 was used.

Files were passed into the kallisto/bustools workflow. For details and scripts on the workflow, see https://github.com/pachterlab/OBP_2025. Because MATQ-seq data were provided as individual FASTQ files corresponding to single bactieral cells, substantial computational overhead was incurred from repeated file opening and ckosign operations. To mitigate this inefficiency and to enable more effective use of the kb-python workflow, a random 8bp sequence was appended to the start of each FASTQ file prior to processing. Files where then concatenated into a signle input FASTQ. This 8bp sequence serves as a cell barcode for downstream analysis.

The resulting count matrix was then loaded into scanpy for basic filtering.

The reads were also passed into the MATQ-seq workflow following the code from https://doi.org/10.1038/s41596-025-01157-5.

When timing was calculated for the MATQ-seq workflow, reference indexing was not included. For kb-python reference index generation was not included.

### Acquisition and preprocessing of BacDrop data

Raw RNA-seq reads from (7) were obtained from GSE141018; specifically GSM5456490/SRR15174657 for the experiment in Fig 4A in (7). Reads were aligned to the https://www.ncbi.nlm.nih.gov/datasets/taxonomy/1328424/ *Klebsiella pneumoniae* BIDMC 35 reference genome.

Following the BacDrop prepreoccesing method, reads were first filtered using UMI-tools to retain only those that contiane both barcode sequences: BC1 (provided in the BacDrop) and BC2 (from the 10X Genomics Chromium Single Cell ATAC barcode whitelist, 737K-cratac-v1.txt.gz, Cell Ranger ATAC v2.2.0). After filtering, UMI, BC1, and BC2 sequences were extracted. Filtered reads were aligned to the reference genome using BWA, gene-level assignments were genetated using featureCounts, and final molecule counts were produced using UMI-tools.

When timing was calculated for the BacDrop workflow, reference indexing was not included. For kb-python reference index generation was not included.

## References

1. Briallen Lobb, Benjamin Jean-Marie Tremblay, Gabriel Moreno-Hagelsieb, and Andrew C Doxey. An assessment of genome annotation coverage across the bacterial tree of life. Microbial Genomics, 6(3):e000341, 2020.

2. Nicolas L Bray, Harold Pimentel, Páll Melsted, and Lior Pachter. Near-optimal probabilistic RNA-seq quantification. Nature biotechnology, 34(5):525–527, 2016.

3. Páll Melsted, A Sina Booeshaghi, Lauren Liu, Fan Gao, Lambda Lu, Kyung Hoi Min, Eduardo da Veiga Beltrame, Kristján Eldjárn Hjörleifsson, Jase Gehring, and Lior Pachter. Modular, efficient and constant-memory single-cell RNA-seq preprocessing. Nature biotechnology, 39(7):813–818, 2021.

4. Delaney K. Sullivan, Kyung Hoi (Joseph) Min, Kristján Eldjárn Hjörleifsson, Laura Luebbert, Guillaume Holley, Lambda Moses, Johan Gustafsson, Nicolas L. Bray, Harold Pimentel, A. Sina Booeshaghi, Páll Melsted, and Lior Pachter. kallisto, bustools and kb-python for quantifying bulk, single-cell and single-nucleus RNA-seq. Nature Protocols, 20(3):587–607, March 2025. ISSN 1750-2799. doi: 10.1038/s41596-024-01057-0. Publisher: Nature Publishing Group.

5. Sydney B. Blattman, Wenyan Jiang, Panos Oikonomou, and Saeed Tavazoie. Prokaryotic single-cell RNA sequencing by in situ combinatorial indexing. Nature Microbiology, 5(10): 1192–1201, October 2020. ISSN 2058-5276. doi: 10.1038/s41564-020-0729-6. Publisher: Nature Publishing Group.

6. Christina Homberger, Fabian Imdahl, Regan J. Hayward, Lars Barquist, Antoine-Emmanuel Saliba, and Jörg Vogel. Transcriptomic profiling of individual bacteria by MATQ-seq. Nature Protocols, pages 1–30, April 2025. ISSN 1750-2799. doi: 10.1038/s41596-025-01157-5. Publisher: Nature Publishing Group.

7. Peijun Ma, Haley M. Amemiya, Lorrie L. He, Shivam J. Gandhi, Robert Nicol, Roby P. Bhattacharyya, Christopher S. Smillie, and Deborah T. Hung. Bacterial droplet-based single-cell RNA-seq reveals antibiotic-associated heterogeneous cellular states. Cell, 186(4):877–891.e14, February 2023. ISSN 0092-8674. doi: 10.1016/j.cell.2023.01.002.

8. Mark D Robinson, Davis J McCarthy, and Gordon K Smyth. edger: a bioconductor package for differential expression analysis of digital gene expression data. bioinformatics, 26(1): 139–140, 2010.

9. Lior Pachter. Differential analysis of genomics count data with edge*. bioRxiv, pages 2026–02, 2026.

10. Matthew Chung, Ricky S. Adkins, John S. A. Mattick, Katie R. Bradwell, Amol C. Shetty, Lisa Sadzewicz, Luke J. Tallon, Claire M. Fraser, David A. Rasko, Anup Mahurkar, and Julie C. Dunning Hotopp. FADU: a Quantification Tool for Prokaryotic Transcriptomic Analyses. mSystems, 6(1):10.1128/msystems.00917–20, January 2021. doi: 10.1128/msystems.00917-20. Publisher: American Society for Microbiology.

11. Laura Luebbert, Delaney K Sullivan, Maria Carilli, Kristján Eldjárn Hjörleifsson, Alexander Viloria Winnett, Tara Chari, and Lior Pachter. Detection of viral sequences at single-cell resolution identifies novel viruses associated with host gene expression changes. Nature Biotechnology, pages 1–10, 2025.

